# Phenotypic plasticity in response to drought stress: Comparisons of domesticated tomato and a wild relative

**DOI:** 10.1101/2023.03.08.531475

**Authors:** Yaniv Lupo, Menachem Moshelion

## Abstract

Phenotypic plasticity is the ability of an organism to undergo reversible behavioral, morphological or physiological changes in response to environmental conditions. Phenotypic plasticity enables plants to cope with uncertain environmental conditions, such as drought. A primary plastic trait is the rate of stomatal response to changes in ambient conditions, which determines the amount of water lost via transpiration, as well as levels of CO_2_ absorption, growth and productivity. Here, we examined the differences between domesticated and wild tomato species and their responses to drought stress. We found that the domesticated tomato had a higher transpiration rate and higher stomatal conductance (g_s_). The domesticated tomato also had greater biomass and greater leaf area under drought conditions, as compared to the wild tomato. Despite the domesticated tomato’s higher transpiration rate and higher g_s_, there was no difference between the photosynthetic rates of the two lines. Moreover, the wild tomato had a higher maximum rate of rubisco activity, which might explain its greater molecular and whole canopy water-use efficiency. The domesticated tomato’s higher transpiration rate and greater leaf area led to its earlier exposure to drought stress, as compared to the wild tomato, which maintained higher levels of soil water, enabling it to maintain steady rates of whole-canopy stomatal conductance (g_sc_) for extended periods. The wild tomato was also more sensitive to the soil water availability and lowered its maximum transpiration rate at a higher soil-water-content level. Our results suggest that the domestication process of tomatoes favored morphological/anatomical performance traits over physiological efficiency.

## Introduction

Plants are immobile organisms and, therefore, are exposed to uncertain and unstable environmental conditions, which can become unfavorable to plant development and even threaten a plant’s survival. Drought is a significant environmental stress that limits crop production and endangers food security (FAO 2020; Dietz et al. 2021). Droughts are expected to become more extensive and intense due to climate change and global warming (Trenberth et al. 2014; Diffenbaugh et al. 2015). Drought affects plants by inhibiting transpiration and photosynthesis, among other processes (Yan et al. 2016). To avoid the adverse effects of drought, plants have evolved different mechanisms to cope with this type of stress (Bacon 2009). One such mechanism is to increase the phenotypic plasticity of stomatal conductance in response to the environment and to engage in more conservative behavior in environments in which the availability of water is unstable (Galkin et al. 2018). However, crop plants have been bred to maximize their productivity during a growing season, while increasing the amount of time that their stomatal are open, despite environmental hazards (Bai & Lindhout 2007). In recent years, it has become increasingly clear that more breeding under unfavorable conditions is essential to minimize the yield gap arising from stressful growing conditions and improve world food security (Cattivelli et al. 2008). Moreover, a better understanding of the mechanisms involved in plant drought response will enable us to breed crops that will perform well in changing environments, to meet the growing demand for agricultural produce (Boyer 1982).

Maintaining high stomatal conductance (g_s_) benefits plants when water is not a limiting factor, since g_s_ is correlated with high yield (Richards 2000; Negin & Moshelion 2016). Although immediate closure of stomata in response to drought helps to ensure survival, maintaining high g_s_ to the point of turgor loss will benefit productivity, but also risks exposing the plant to more severe stress and endangering its survival (Sade et al. 2012). Therefore, the ideal, plastic stress response is high g_s_ under well-watered conditions and low g_s_ under drought conditions (Negin & Moshelion 2016). Moreover, g_s_ is continually regulated to maximize the momentary water-use efficiency (WUE) in response to the dynamic soil-atmosphere conditions (Gosa et al. 2019). A high WUE is desirable for agriculture, as it allows crops to produce more yield using less water. The same goes for wild plants, as high WUE benefits competition and survival under limited-water conditions.

Conventional plant-breeding processes are based on the hybridization of parents and phenotypic selection of offspring and are relatively slow. An average breeding program for an annual crop can take 10–12 years (Kumar et al. 2015; Spindel & McCouch 2016) depending on factors such as the target environment, the availability of genetic variation and the heritability of the trait of interest. In recent decades, the development of technologies associated with molecular markers and genomic selection has provided new tools that have enhanced classical breeding processes and made breeding for simple and complex traits more efficient (Spindel et al. 2015; Bhat et al. 2016). Today, the technical challenge of genomic selection lies in the reliability of the available phenotypic data. The gap between the genotypic data and the phenotypic data available to breeders creates a situation known as the genotype–phenotype gap. The genotype–phenotype gap is even more complex when it involves interactions with the environment and specific interactions with a water-limited environment. Therefore, to harness the full benefits of new genetic technologies, we must apply them with high-throughput phenotyping under different environmental conditions. Functional physiological phenotyping (FPP) is a physiology-based, high-throughput, non-destructive and non-invasive phenotyping technique that continually measures the plant and ambient conditions (i.e., soil and atmosphere). FPP enables the detection of small changes in specific physiological traits (e.g., g_s_) associated with environmental changes, in general, and stress conditions, in particular (Gosa et al. 2019).

In this work, we investigated whether the phenotypic plasticity of the stomatal response to drought stress has been preserved in a crop plant after centuries of domestication and selective breeding. For this investigation, we used a domesticated tomato (*Solanum lycopersicum* cv. M82) and its wild relative (*Solanum pennellii*). We worked with these species for four main reasons. First, tomato is an important crop plant. Second, this choice allowed us to limit genetic variation as both lines are self-compatible. Third, introgression lines of these two species are available for future research (Eshed & Zamir 1995). Finally, *S. pennellii* has adapted to its native environment, the arid regions of the Andes in South America, as can be seen in its physiological drought-response patterns (Bolger et al. 2014).

We used the FPP system to examine many plants over short (i.e., hours) and long (i.e., weeks) periods (Halperin et al. 2017; Dalal et al. 2020). With this system, we were able to simultaneously analyze continuous data during periods of optimal irrigation and drought. We hypothesized that M82 would have higher absolute levels of physiological and morphological traits related to plant water balance (e.g., photosynthesis, stomatal conductance, growth rate and WUE) and would be more efficient, but also more sensitive to drought than *S. pennellii*.

## Materials and Methods

### Plant material

For this study, we used a domesticated tomato (*Solanum lycopersicum* cv. M82) and its wild relative (*Solanum pennellii*; referred to as *Pennellii*). Seeds from each plant line were sprouted in a commercial germination plate and were transferred to the FPP system after four weeks.

### Experimental setup

The experiment was conducted in a controlled greenhouse at the Faculty of Agriculture, Food and Environment in Rehovot, Israel (iCORE functional-phenotyping greenhouse) during the spring of 2017. The iCORE polycarbonate greenhouse includes cooling pads and a heater, which were activated as needed to maintain a minimum temperature of 14ºC. The light in the greenhouse is natural sunlight. The continuous photosynthetically active radiation (PAR) and vapor pressure deficit (VPD) measured during the experimental period are presented in Fig. 3a. Climate conditions were continuously monitored (recorded every 3 min) by a weather station, which is a part of the Plantarray® functional phenotyping platform in the greenhouse. Plants were cut at the bottom of the stem at the end of the experiment for the measurement of fresh and dry weight (*n* = 18 for M82 and *n* = 31 for Pennellii). Agronomic and canopy WUE were calculated from the linear regression line of shoot dry or fresh weight taken during the plants’ vegetative growth stage (days 13/05-06/06), respectively, and the total transpiration. Leaf area was measured at the end of a preliminary experiment under similar environmental and stress conditions, as described in this experiment.

### Functional-phenotyping platform

The functional-phenotyping platform (FPP; Plantarray®, Plant-Ditech Ltd., Yavne, Israel) was used to monitor plant growth and water balance through controlled tracking and measurement of the transpiration and biomass gain of each plant throughout the growing period (Fig. 1). The system uses highly sensitive, temperature-compensated load cells, which serve as weighing lysimeters. Each lysimeter unit was connected to its own controller, which collected data and controlled the irrigation of each 3.8-L pot filled with growing media and planted with one plant (described in greater detail below). Together with different sensors incorporated into the system, each plant’s soil–plant–atmosphere continuum was measured independently and simultaneously throughout the experimental period. The Plantarray® system recorded the weights of the pots and the environmental information that was registered by the sensors every 3 min. The data collection could be viewed in real-time through the online web-based software SPAC-analytics (Plant-Ditech). The setup and measurement techniques of the system are described thoroughly in Halperin et al. (2017) and Dalal et al. (2020).

**Figure 1.**
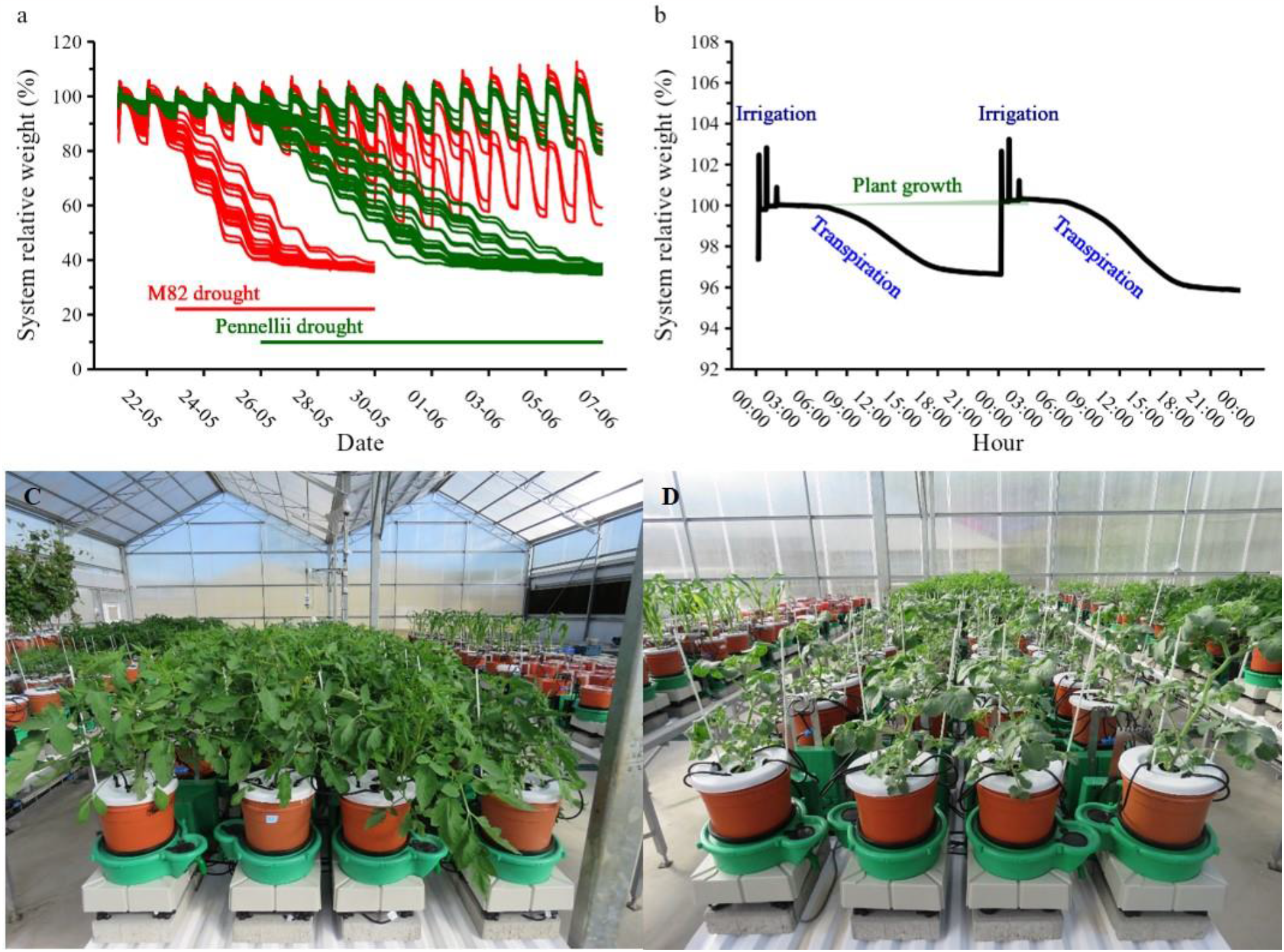
Raw data from the FPP system. (A) Raw data regarding the system’s relative weight (plant + pot + container) throughout the experiment. Each line represents one plant; red = M82, green = Pennellii. (B) Representative two days of the system under the control condition showing the daily loss of weight from transpiration, three irrigation peaks during the night and the plant growth rate calculated from the increase in the system weight. (C) M82 plants on the FPP on May 22 ^nd^. (D) Pennellii plants on the FPP on May 22^nd^.

The pots were filled with a commercial growth medium (Bental 11, Tuff Merom Golan Ltd.), which will be referred to as *soil*. Prior to the filling of the pots, the soil was mixed for several hours with excess water in a commercial-size mixer to achieve soil saturation (∼80% volumetric water content). Each pot was put into a plastic container through a hole in its cover. Evaporation from the containers and from the soil was prevented by a plastic cover with a hole in the middle to allow the plant stem to emerge. Prior to the experiment, all load cells were calibrated with 1-kg and 5-kg weights and randomly examined for reading accuracy.

Each plant was irrigated by four on-surface 4-L/h drippers to ensure uniform water distribution in the pots at the end of the irrigation event and prior to free drainage. Plants were irrigated in three consecutive cycles of irrigation and drainage between 24:00 and 02:00. Fertilizer (Haifa Group, Poly-FeedTM, N-P-K 17-10-27 + ME) was applied with the irrigation water. After the pots had finished draining, the daily pre-dawn pot weight was determined as the average weight between 04:00 and 04:30. Additional water was provided during the irrigation event and was stored in the container, on average an additional 250 mL. The amount of additional water was managed by a drainage orifice in the container side wall through which the excess water drained out. The reservoir water ensured that water would be fully available to the well-irrigated plants throughout the following day without supplemental irrigation. Without additional irrigation during the day, it is possible to ensure a monotonic decrease in pot weight over two consecutive days. This method enabled us to apply the data-analysis algorithm SPAC-analytics. Reaching an a priori-determined water level at drainage completion during the night enabled us to determine the daily plant weight gain (Fig. 1b, 3c).

Transpiration was calculated as the difference between the weight just before dawn (when SWC was at field capacity) and the weight in the evening (after sunset). Transpiration was later normalized to the plants’ weight (E_c_) or to the plants’ weight and the VPD (g_sc_). The experimental study included two treatments: surplus irrigation and drought. The drought treatment was induced by stopping irrigation (for more details, see Halperin et al. 2017). We were also able to use our real-time monitoring system to evaluate each plant’s stress level and decide when to start and stop the drought treatment. Both M82 and Pennellii were grown on the FPP. The drought treatment started when the plans reached 650 (M82; 23/05) and 230 (Pennellii; 26/05) mL of daily transpiration. The drought treatments continued until the treated plants reached 10% SWC (7 days for M82 and 12 days for Pennellii; Fig. 3f).

### Gas-exchange measurements

Gas-exchange measurements were taken with the LI-6400XT portable gas-exchange system equipped with a 6-cm^2^ aperture standard leaf cuvette (LI-COR, Lincoln, NE, USA; Fig. 5). Samples were taken from young, fully expanded, sun-exposed leaves from the top of the plant. The plants used for the gas-exchange measurements were grown next to the plants on the FPP and under the same conditions as the control plants. Measurements were taken under saturating light (1200 μmol m^-2^ s^-1^; blue light was set to 10% of the photosynthetically active photon) with 400 μmol mol^-1^ CO_2_ surrounding the leaf flux density. The leaf-to-air VPD was kept between 1.5 and 2.5 kPa during all measurements. The leaf temperature for all measurements was about 25°C. Measurements for A–Ci curves were taken under the same conditions described above (Fig. 6). The CO_2_ surrounding the leaf flux density ranged from 50 μmol mol^-1^ to 1600 μmol mol^-1^. A–Ci curves and the maximum rate of carbon fixation (V_cmax_) were calculated with the Plantecophys R Package, as described by (Duursma 2015).

### Data analysis

Data from the FPP system were analyzed with the data-analysis algorithm SPAC-analytics. The data set included daily data (one value per day per plant) and momentary data acquired every 3 min (480 values per day per plant). The absence of overlapping confidence intervals indicates a significant difference at *P* < 0.05 (Fig. 4a, b). Other data analyses and data visualization were performed using R (4.2.1). Midday averages were fitted to a smooth curve using locally weighted polynomial regression (Fig. 3b, d and e). Means deemed significantly different at *P* < 0.05 were compared using Student’s *t*-test or ANOVA followed by Tukey-Kramer’s HSD (noted in the figure legends). Analysis of covariance (ANCOVA) was used to compare the slopes in Figs. 2b and 2d.

**Figure 2.**
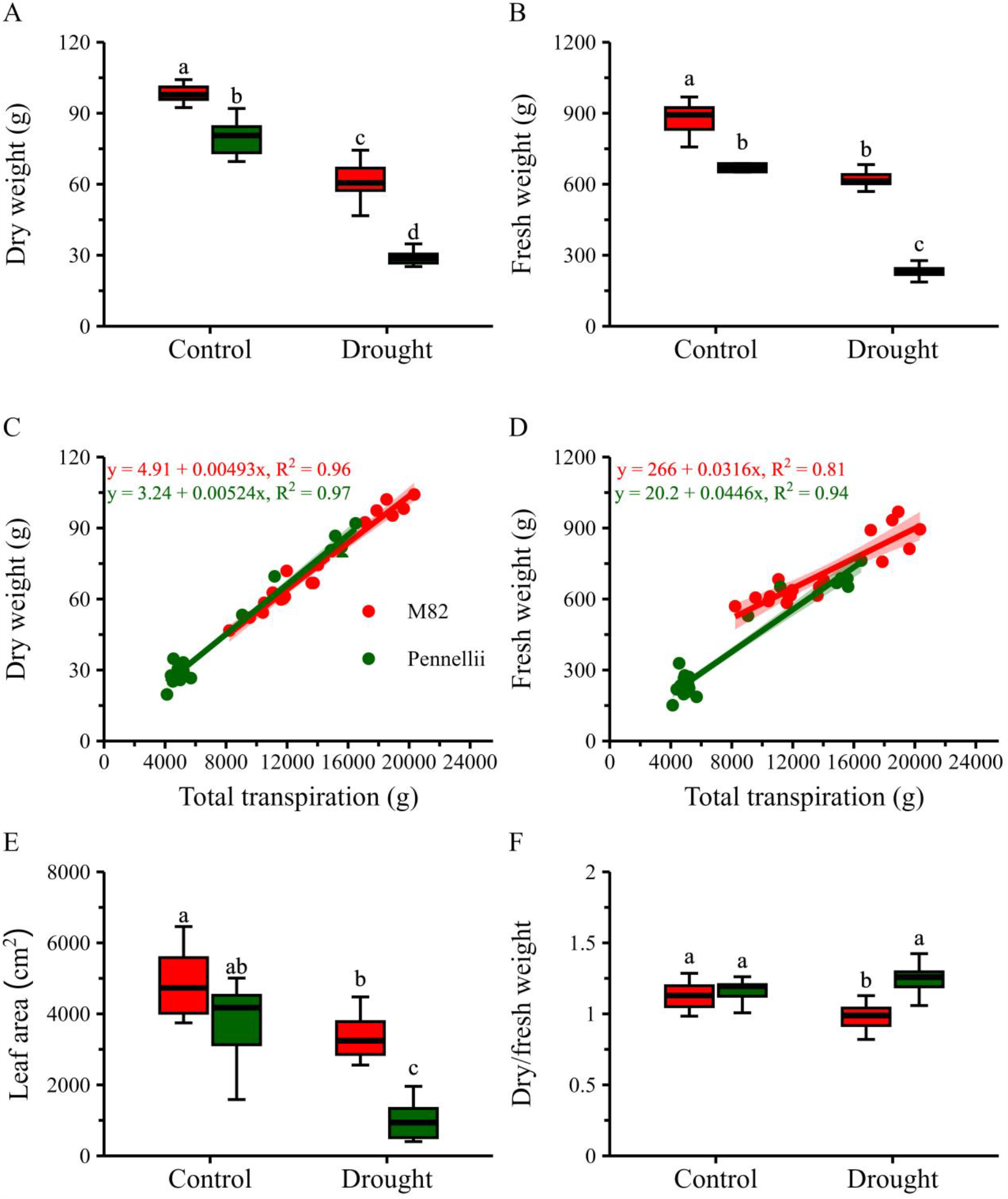
Morphological parameters and water-use efficiency (WUE) at the end of the experiment. (A) Plant dry weight. (B) Plant fresh weight. (C) Agronomic WUE (dry weight vs. total transpiration). (D) Canopy WUE. (fresh weight vs. total transpiration) (E) Leaf area (LA). (F) Dry-to-fresh weight ratio; red = M82, green = Pennellii. Different letters represent significant differences between species and treatments, according to Tukey’s HSD, *P* < 0.05. *n*_weight_ = 6 (M82 control), 12 (M82 drought), 7 (Pennellii control) and 24 (Pennellii drought). *n*_LA_ = 8 (M82 control), 12 (M82 drought), 6 (Pennellii control) and 17 (Pennellii drought).

**Figure 3.**
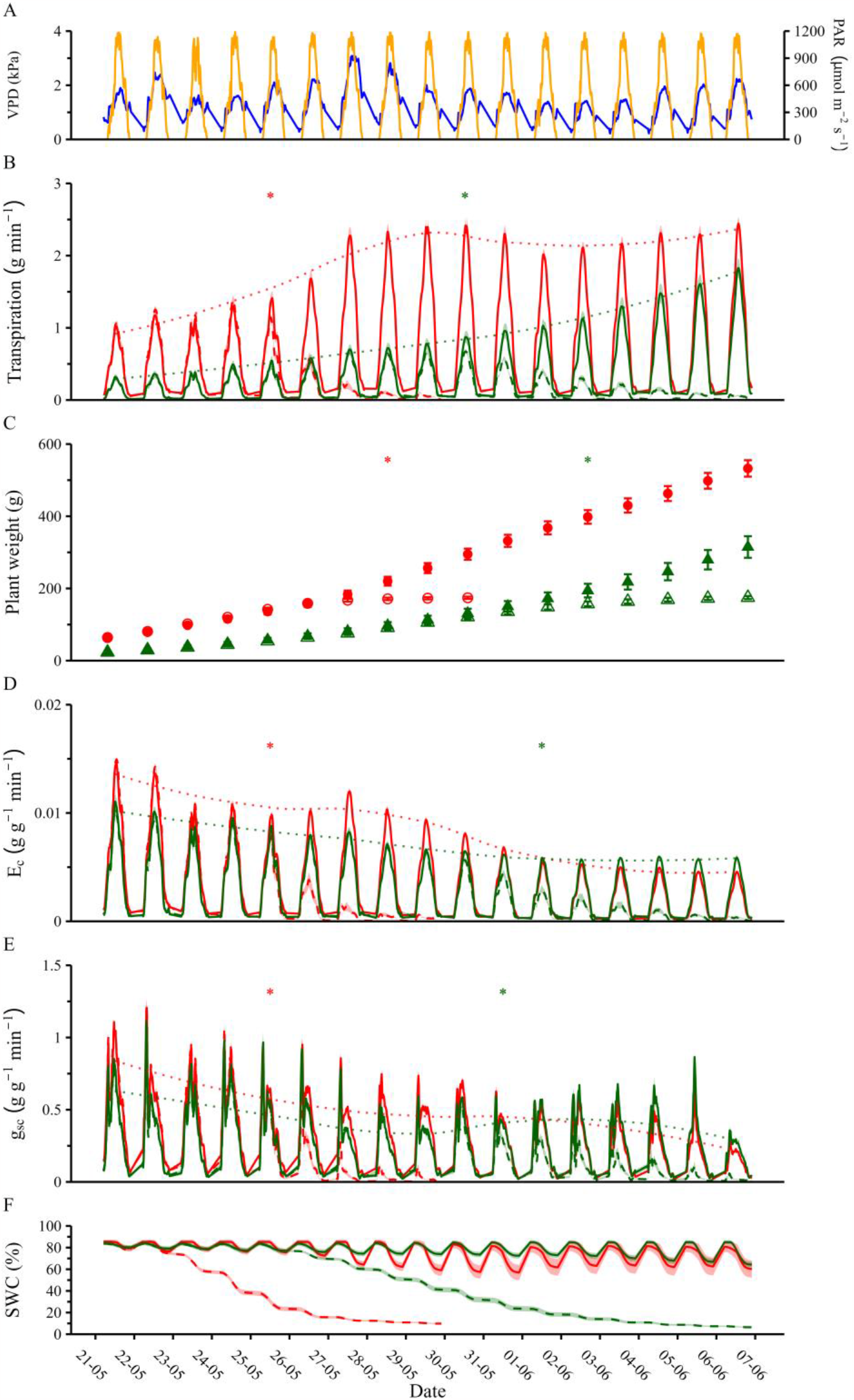
Soil-plant-atmosphere-continuum (SPAC) data. (A) Vapor pressure deficit (VPD; blue line) and photosynthetic active radiation (PAR; orange line) over time. (B) Absolute transpiration rate (whole-plant water-loss rate, continuous plot) and midday average (dotted line) over time. (C) Calculated plant weight over time; circles = M82, triangles = Pennellii. (D) Transpiration rate normalized to plant weight (E_c_, continuous plot) and midday average (dotted line) over time. (E) Whole-canopy stomatal conductance (g_sc_, continuous plot) and midday average (dotted line) over time. (F) Calculated soil volumetric water content (SWC) over time. (B, D–F) Red = M82, green = Pennellii; solid lines/full symbols = control, dashed lines/empty symbols = drought. Asterisks represent the first day on which significant differences between treatments within each species were observed, according to Student’s *t*-test, *P* < 0.05. *n* = 7 (M82 control), 23 (M82 drought), 7 (Pennellii control) and 24 (Pennellii drought).

**Figure 4.**
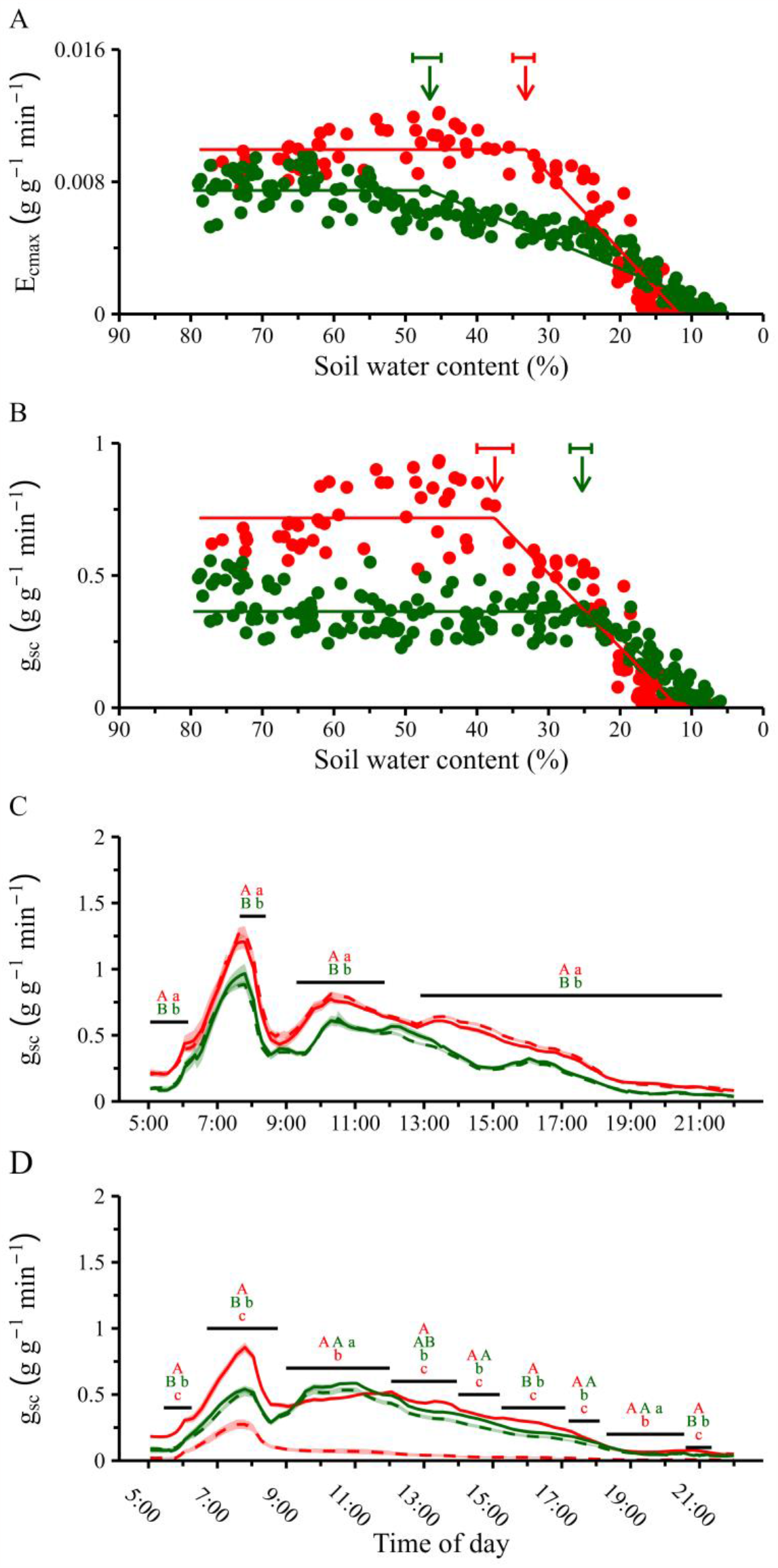
Piecewise curve fit and diurnal canopy stomatal conductance at different soil water content (SWC) levels. (A) Piecewise curve fit of midday E_c_ in response to SWC. (B) Piecewise curve fit of midday g_sc_ in response to SWC. (C) Diurnal g_sc_ under well-irrigated conditions (SWC = ∼80%). (D) Diurnal g_sc_ 5 days after drought started (SWC at 5 am was 16% and 39% for M82 and Pennellii, respectively). Red = M82, green = Pennellii; solid lines = control, dashed lines = drought. Arrows mark the point at which soil water is restricted from supplying midday transpiration needs (Ɵ_crit_). Bars represent upper and lower CI for Ɵ_crit_. Different letters represent significant differences between species and treatments (uppercase represents control and lowercase represents drought), according to two-way ANOVA, *P* < 0.05. *n* = 7 (M82 control), 23 (M82 drought), 7 (Pennellii control) and 24 (Pennellii drought).

## Results

At the end of the experiment, M82 had higher fresh and dry shoot weights under the control and drought treatments, as compared to Pennellii (Fig. 2a, c). Both species had lower fresh and dry weights under the drought treatment than under the control treatment. M82 also had greater leaf area than Pennellii under the drought treatment. Both species had smaller leaf areas under the drought treatment than under the control condition (Fig. 2e). There were no significant differences in the agronomic WUE of the two species (0.52% for Pennellii as compared to 0.49% for M82; slope in Fig. 2b). Pennellii had higher canopy WUE (4.46%, as compared to M82’s 3.16%; slope in Fig. 2d). M82 had a lower dry/fresh weight ratio under the drought treatment, as compared to the control and as compared to Pennellii (Fig. 2f).

Comparative functional-phenotyping screening of whole-plant water-balance regulation was conducted under similar ambient VPD and PAR conditions and those parameters were measured continuously throughout the experiment (Fig. 3). Under the control condition, M82’s absolute transpiration rate (non-normalized) was higher than Pennellii’s throughout the period of the study (Fig. 3b). M82 and Pennellii exhibited lower transpiration rates (compared to the control) at 3 and 5 days, respectively, after drought treatment was initiated (Fig. 3b). Under well-irrigated conditions, both M82 and Pennellii exhibited an increase in transpiration rate from day to day. However, throughout the experiment, M82 showed a faster rate of increase due to the fact that it was gaining mass more quickly (Fig. 3c). In response to the drought treatment, M82 stopped growing after 6 days (28/05); whereas Pennellii stopped growing after 12 days (06/06). When total transpiration was normalized to plant weight (E_c_), the differences between M82 and Pennellii were reduced, yet M82 still had a higher E_c_ under the control condition (Fig. 3d). Moreover, both M82 and Pennellii exhibited decreasing E_c_ over the course of the experiment. A similar pattern was revealed also for the canopy stomatal conductance (g_sc_; Fig. 3e).

Plotting the midday E_c_ (E_cmax_) against the SWC revealed the transpiration response to the reduction in SWC (Fig. 4a). We found that M82 maintained higher E_max_ through lower SWC, reaching the lower physiological drought point (θ_crit_; determined as the point at which soil water limits the transpiration needs) at 33.2% and then decreasing sharply (slope = 0.05). Pennellii, on the other hand, exhibited a much stronger water-conservative response of significantly higher θ_crit_ of SWC = 46.6%, followed by a more moderate decrease (slope = 0.02). Plotting M82’s midday g_sc_ response pattern to SWC revealed a similar response pattern to E_c_. Nevertheless, Pennellii’s g_sc_ responded differently than its E_c_, suggesting different regulation of the whole-canopy conductance under drought. To examine the g_sc_ pattern at a higher resolution, we looked at momentary g_sc_ throughout the day (Fig. 4c, d). Plants were compared 1 day before the drought was initiated (22 May and 25 May for M82 and Pennellii, respectively) and 5 days after irrigation was stopped. The atmospheric conditions were similar on those days. Under well-watered conditions, M82 had a higher g_sc_ than Pennellii throughout the day (Fig. 4c). In terms of transpiration, Pennellii was more sensitive to drought than M82 was (higher θ_crit_), which enabled it to keep more available soil water for a more extended period. This allowed it to maintain a higher g_sc_ (as compared to M82) after 5 days of drought, as could be detected from the fact that there were no differences between the Pennellii control and drought treatments throughout most of the day (Fig. 4d). At the same time, throughout the day, the g_sc_ of drought-treated M82 was lower than that of the control M82 and that of the drought-treated Pennellii (Fig. 4d).

Gas-exchange measurements at the single-leaf level revealed no differences in the CO_2_ assimilation rates (A_n_) of M82 and Pennellii under well-irrigated conditions (Fig. 5a). M82 had higher g_s_ and E than Pennellii (Fig. 5b, c). This meant that Pennellii had higher iWUE under control conditions, as compared to M82 (Fig. 5d). Moreover, Pennellii also had a higher maximum carboxylation rate (V_cmax_) than M82 (Fig. 6b). There were no significant differences between the two species’ maximum rates of electron transport (J_max_; Fig. 6c).

**Figure 5.**
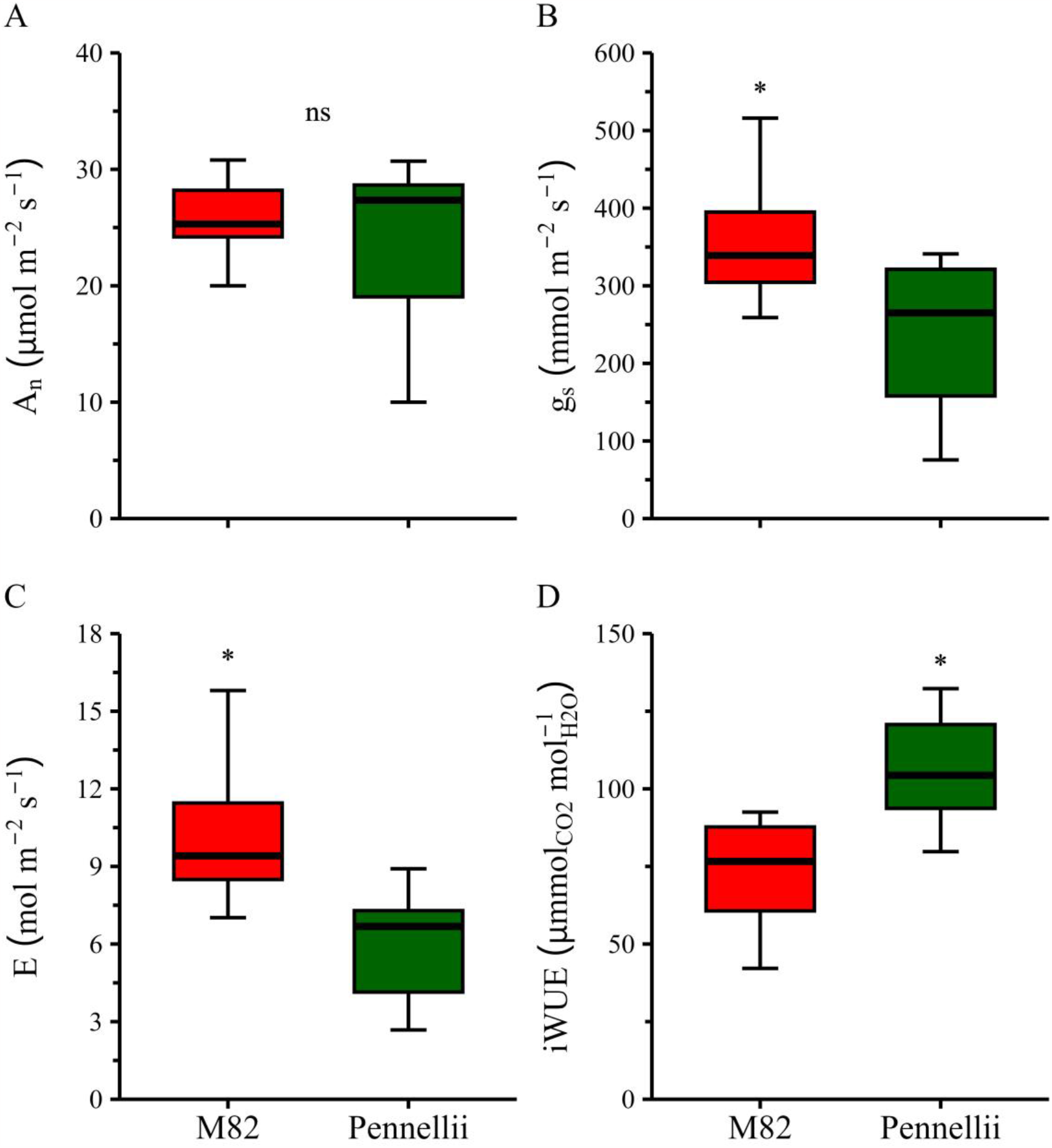
Gas exchange of well-watered plants of the two species. (A) Photosynthetic assimilation rate (A_n_). (B) Stomatal conductance (g_s_). (C) Transpiration rate (E). (D) Intrinsic water-use efficiency (iWUE; calculated as A_n_/g_s_). Red = M82; green = Pennellii. Asterisks represent significant differences according to Student’s *t*-test, *P* < 0.05. *n* = 11 (M82) and 10 (Pennellii).

**Figure 6.**
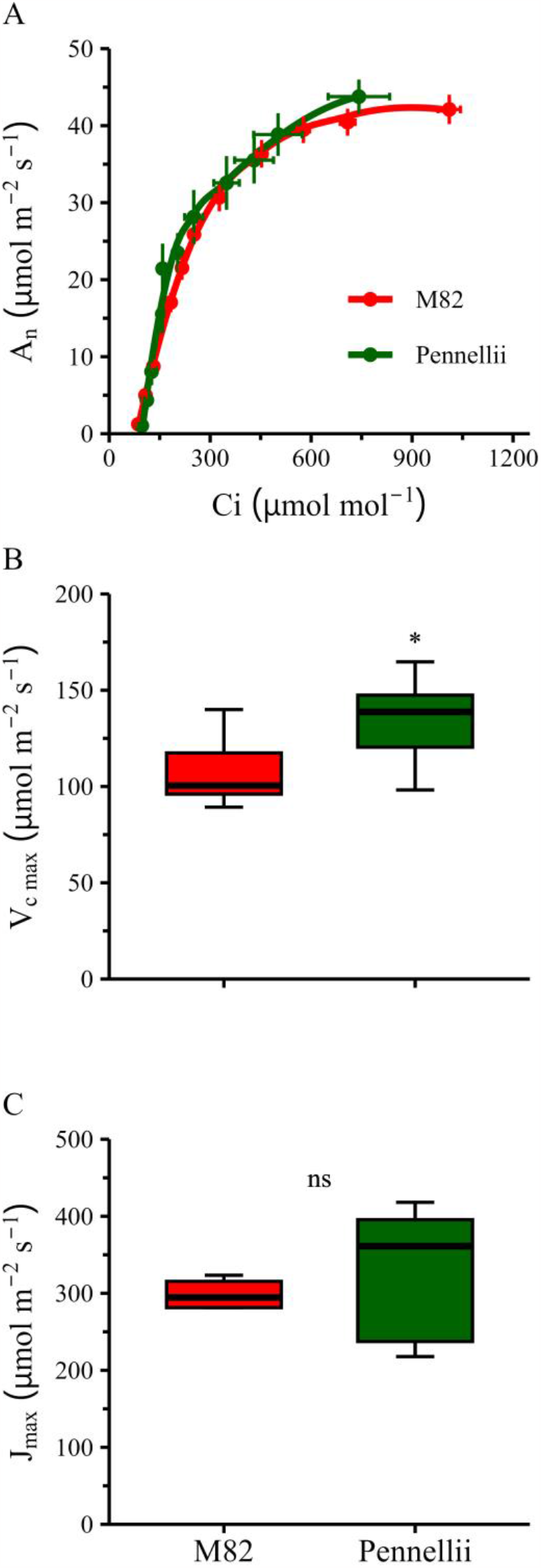
CO_2_ response curves of the two species. (A) A–Ci curves. (B) Maximum rate of carbon fixation (V_cmax_). (C) Maximum rate of electron transport (J_max_). Red = M82; green = Pennellii. An asterisk represents a significant difference according to Student’s *t*-test, *P* < 0.05. *n* = 11 (M82) and 10 (Pennellii).

## Discussion

In this study, we evaluated the water-balance responses of cultivated and wild tomato species to drought. The domesticated species (M82) had greater biomass, grew more quickly and exhibited greater stomatal conductance and greater total transpiration than the wild species (Pennellii). However, there were no differences in the species’ photosynthetic rates per unit leaf area, which resulted in Pennellii’s higher WUE (canopy and intrinsic, Fig. 2 and Fig. 5, respectively). M82 was also less sensitive to declining SWC than Pennellii, as it maintained a high level of transpiration at low SWC levels. M82’s higher total transpiration, combined with its slower response to reduced SWC, led to faster and more intense drought stress for M82, as compared to Pennellii.

These findings are congruent with our hypothesis that cultivated plants are more canalized toward production and, therefore, exhibit a less plastic stomatal response to the environment, resulting in a risk-taking response to drought. However, unlike what we hypothesized, the cultivation process does not appear to have resulted in more efficient behavior, but does appear to have increased the plant size and, in doing so, increased the total transpiration and photosynthesis, to support higher yields.

### Domestication led to a high-transpiring and more drought-susceptible plant

Domestication and breeding of tomato led to changes in a wide range of traits following a process referred to as the domestication syndrome (Harlan 1992; Bai & Lindhout 2007). One of the domestication traits is the increase in plant size to support high yields. In this work, we demonstrate that M82 has not only greater plant mass, but also higher total and normalized (to biomass) rates of transpiration due to its higher g_sc_, as compared to Pennellii. High g_sc_ and a high transpiration rate can be beneficial when environmental conditions are not limiting, as they allow for more gas exchange and CO_2_ fixation. Yet, higher transpiration often has a tradeoff with efficiency (Dalal et al. 2017). Indeed, M82’s higher transpiration resulted in lower WUE (Figs. 2 and 5). Examination of the relationship between E_cmax_ and SWC shows that Pennellii reduced its transpiration at a higher SWC (Ɵ_crit_) than M82, thereby maintaining a higher SWC for a more extended period of time (Figs. 3f and 4a). Interestingly, this behavior enabled Pennellii to keep growing for a longer period under drought (Fig. 3c) and to maintain constant whole-canopy conductance over those longer periods (Fig. 4b). In contrast, M82’s higher g_sc_ and higher total transpiration resulted in a faster reduction in SWC under terminal drought, which reduced g_sc_ and plant growth almost immediately after the Ɵ_crit_ of E_cmax_ was reached. M82’s higher E_cmax_ combined with its late response to drought led it to experience drought stress faster than Pennellii, but also allowed it to be more productive until that lower Ɵ_crit_ point was reached.

At the leaf level, even though M82 had higher g_s_ and E than Pennellii, there were no differences in the A_n_ levels of the two species (Fig. 5a). This combination of lower transpiration and a similar CO_2_ fixation rate led to Pennellii’s higher iWUE (Fig. 5b), which was in agreement with its higher canopy WUE (Fig. 2b). Moreover, Pennellii also exhibited more efficient CO_2_ fixation (V_cmax_; Fig. 6b), which explains how it can fix more CO_2_ at a lower g_s_ (Walker et al. 2014). This explains Pennellii’s higher dry/fresh shoot weight ratio (Fig. 2f). Therefore, Pennellii is more adaptable to dry conditions, as it can fix more CO_2_ and produce more biomass with less water lost to transpiration.

Taking all the above into consideration, we can say that Pennellii may have developed a more conservative water-protective drought adaptation than M82, which involves transpiring less and responding more quickly to the depletion of water from the soil, as could be expected from a wild species evolving toward “survivability-enhancing” behavior (Dalal et al. 2017). Pennellii is native to an arid region, in which it probably adapted to become drought-tolerant, in order to survive. M82, on the other hand, is a domesticated plant bred to produce high yields under optimal conditions, which may result in it being less able to survive under prolonged drought. Yet, the question remains, why weren’t these beneficial ecological traits of Pennellii maintained through the breeding process?

### Domestication for yield led to a higher transpiring, but less plastic and less efficient plant

Domestication and cultivation started long before biochemical techniques were available. Plants were selected for simple morphological/agronomical traits, such as size and yield. In the process of breeding M82 for rapid growth and high yield under optimal water conditions, its WUE was inadvertently reduced. The physiological definition of molecular WUE (units of CO_2_ gained in photosynthesis per units of H_2_O lost in transpiration) measures efficiency, but not productivity. Indeed, M82 is less efficient and had lower iWUE (A_n_/g_s_), but is more productive in terms of biomass gain and is reported to have higher yields in the field (Eshed & Zamir 1995), indicating a tradeoff between efficiency and productivity.

These results match what has been found in other crops. For example, a comparison of cultivated wheat with its wild progenitors showed that the area of individual leaves and the total leaf area of seedlings increased with the shift from wild to cultivated forms, but that increase was coupled with a progressive reduction in the rate of photosynthesis per unit leaf area (Evans & Dunstone 1970). Moreover, in the last century, there has been no change in the rate of photosynthesis in cereals, yet total photosynthesis has increased as a result of an increase in total leaf area, the daily duration of photosynthesis or leaf area, but not due to any direct improvement of photosynthesis efficiency (Richards 2000). However, these robust biochemical traits seem to have been canalized in modern crops, leaving the question of their native plasticity unanswered.

Plasticity refers to the ability of a plant to dynamically change its phenotype in response to changes in its environment. Adaptive phenotypic plasticity benefits plants by enabling them to produce better phenotype–environment matches under different environmental conditions (DeWitt et al. 1998).

Stomatal movement is a highly plastic trait that allows the plant to optimize its risk/production response to a dynamically changing environment. High plasticity for this trait is beneficial as it helps the plant to avoid the adverse effects of drought as it rapidly responds to small changes in the SWC, yet it carries the penalty of reduced photosynthesis and production. M82 exhibited much less plasticity in this trait, compared with Pennellii, as it maintained higher transpiration, only reducing its E_max_ at a relatively low SWC, leading to rapid exposure to severe stress (Fig. 4a). In Pennellii, on the other hand, this trait was more plastic. Pennellii exhibited lowered E_max_ under higher SWC, which enabled it to reserve more water and gave it more time to adjust to the drought. At that point, the plants maintained their rate of growth (Fig. 3c) and exhibited a slower reduction in g_sc_ (Fig. 3e), which is an additional aspect of Pennellii’s plastic behavior, seen in its lower Ɵ_crit_ of g_sc_ (Fig. 4b).

An additional plastic trait is the daily pattern of stomatal response. Interestingly, under well-irrigated conditions, both lines exhibited a morning peak of high g_sc_ when VPD was relatively low and PAR was sufficient, thus maintaining optimal behavior. This peak is referred to in the literature as the “golden hour” and correlates with tomato yield in the field (Gosa et al. 2022). Pennellii’s rapid E_c_ response to lowered SWC actually enabled it to maintain the golden-hour peak plasticity for longer periods under drought, as compared with M82 (Fig. 4c, d). This behavioral tradeoff is very interesting as it enabled Pennellii to reduce its risk through fast closure of stomata during the part of the day when VPD is higher and although it had lower g_sc_, it could be maintained for a longer time, which probably improves the plant’s survival chances. Yet, on the other hand, this activity results in smaller plants at the end of the drought period (Fig. 2). Interestingly, under well-irrigated conditions, this daily plasticity peak was maintained in M82 at an even higher level. The availability of introgression lines between M82 and Pennellii can be used in future trials to evaluate the golden-hour peak effect on WUE and productivity.

Our results suggest that the cultivation process canalized (fixed) plastic traits, such as high stomatal conductance, low responsiveness to drought and the golden-hour peak, thereby directly improving yield under well-watered conditions (i.e., the conditions used in the breeding process). Moreover, we suggest that breeding had more of an impact on the rate of growth than on biochemical (photosynthetic) activity traits. At the same time, it reduced a “protective” plasticity response to the environment, while maintaining productive plasticity such as the golden-hour peak, which enables dynamic WUE over the course of the day (Fig. 4a, c). Therefore, we conclude that breeding increased M82’s biomass and, in doing so, also increased its total transpiration and photosynthesis, resulting in increased yield, but at the cost of a decreased ability to sense and respond to environmental changes, which put it under greater risk in stressful conditions.

### Functional phenotyping under stress: Should we normalize when evaluating plant performance?

Transpiration measurements are essential for evaluating plant water status and plants’ responses to their environment, particularly under drought. Should we use normalized or absolute transpiration when comparing plants? Should we use leaf or whole-plant measurements to estimate drought response? Normalization of transpiration helps when comparing different plants observed at different times. However, normalization can be misleading and even inaccurate in some cases, such as when the assumptions used are not tested and proved for every examined line. For example, a broadly accepted assumption for gas-exchange calculations used in many studies over the past decades is that the relative humidity inside the substomatal cavities is near-saturated (Farquhar & Raschke 1978). However, in *Eucalyptus pauciflora*, the relative humidity inside the intercellular air spaces at midday was found to be 90% (Canny & Huang 2006). A recent study by Farquhar (Wong et al. 2022) also contradicted the original assumption by showing that the RH inside the substomatal cavities decreased to 80% when the air humidity was reduced. Moreover, an improved method for calculating gas exchange was recently suggested (by the author of the original theory), which also considers the cuticular conductance to water (Márquez et al. 2021). However, this improved method is still valid only for leaf-level measurements. The selected leaves for this kind of measurement are usually young and sun-exposed and do not represent the entire plant canopy. As the plant grows, leaves mature and become less productive and the basal leaves are shaded, resulting in reduced canopy conductance. Therefore, using non-normalized whole plant measurements (e.g., absolute transpiration) is essential to better understand the whole plant’s response to the environment based on values that need no interpretation (e.g., mL water transpired per plant per day). Moreover, absolute transpiration better demonstrates the whole plant’s adaptation to the environment by showing the actual soil–plant water balance, which can be also interpreted to determine the exact irrigation amount.

For example, in this work, M82 had much higher absolute transpiration than Pennellii when there was an adequate supply of water, which led those plants to reach their pot capacity (∼1200 mL) on May 30^th^ (Fig. 3b). However, normalizing the transpiration to plant weight (E_c_) or to plant weight and VPD (g_sc_) reduced the size of the differences between the species (Fig. 3d, e), despite the big difference in their actual water loss.

Measuring the entire plant canopy allows us to avoid the need to decide which leaf to choose for manual gas-exchange measurements, as the system measures the whole plant, including old and shaded leaves. The actual whole-plant g_sc_ exhibits a declining trend (Fig. 3e), resulting from the fact that many of the older leaves maintain their size and weight as their performance decreases and lower leaves becomes shaded. Moreover, measuring the entire canopy with no physical interference (e.g., clipping chamber, wind blowing or other effects that the manual gas-exchange apparatus impose on a leaf) that modifies the boundary layer or any assumptions regarding the abaxial/adaxial stomatal density provides an accurate measure of canopy stomatal conductance, as recently reported for the Plantarray system (Jaramillo Roman et al. 2021).

Therefore, in plant-breeding programs in which many cultivars are screened in parallel and compared based on their performance, we suggest that whole-plant non-normalized behavior will be better represented by the absolute water consumption, which presents the field scenario and helps the breeder to see the actual risks and behavioral response of the different lines. In terms of WUE, breeding for improved canopy-level WUE is essential in the face of a changing climate (Hatfield & Dold 2019). However, if we are aiming to improve crop plant productivity, the WUE must be integrated with yield production and cannot serve as a stand-alone trait. Here, we show how high-throughput phenotyping of a crop’s wild relative may gain potential for improving canopy WUE, under unfavorable environmental conditions.

## Conclusions

The domestication and breeding of M82 led to significant increases in mass and absolute transpiration, resulting in higher absolute photosynthesis and increased yields under non-water-limited conditions. However, under drought stress, M82 exhibits a more rapid reduction in soil water content than Pennellii does. Our study revealed that the domestication and breeding process primarily impacted M82’s morphological and physiological traits rather than its biochemical traits. It appears that high-yielding plants were favored during the breeding process, with less emphasis placed on WUE, likely due to the difficulty of measuring that trait. Our findings suggest that a tradeoff exists between a plant’s productivity and its adaptability to different environments and that during crop domestication this balance can shift toward productivity at the expense of plasticity. With the increasing scarcity and cost of agricultural resources such as water and fertilizers, future breeding programs should prioritize plants’ physiological adaptation to changing environments in which resources may be limited.

## Author contributions

Y.L. and M.M. conceived the idea presented in this work. Y.L. conducted the experiment, and M.M. supervised the findings. Y.L. and M.M. discussed the results, made contributions to the final manuscript, and both wrote the paper.

## Acknowledgements

The work was funded by the Israel Innovation Authority (grants No. 0002122 and 001897).

## Data availability statement

The original data presented in the study are included in the article and supplementary material. Further inquiries can be directed to the corresponding author.

